# Bayesian transfer in a complex spatial localisation task

**DOI:** 10.1101/716431

**Authors:** Reneta Kiryakova, Stacey Aston, Ulrik Beierholm, Marko Nardini

## Abstract

Prior knowledge can help observers in various situations. Adults can simultaneously learn two location priors and integrate these with sensory information to locate hidden objects. Importantly, observers weight prior and sensory (likelihood) information differently depending on their respective reliabilities, in line with principles of Bayesian inference. Yet, there is limited evidence that observers actually perform Bayesian inference, rather than a heuristic, such as forming a look-up table. To distinguish these possibilities, we ask whether previously-learnt priors will be immediately integrated with a new, untrained likelihood. If observers use Bayesian principles, they should immediately put less weight on the new, less reliable, likelihood (“Bayesian transfer”). In an initial experiment, observers estimated the position of a hidden target, drawn from one of two distinct distributions, using sensory and prior information. The sensory cue consisted of dots drawn from a Gaussian distribution centred on the true location with either low, medium, or high variance; the latter introduced after block three of five to test for evidence of Bayesian transfer. Observers did not weight the cue (relative to the prior) significantly less in the high compared to medium variance condition, counter to Bayesian predictions. However, when explicitly informed of the different prior variabilities, observers placed less weight on the new high variance likelihood (“Bayesian transfer”), yet substantially diverged from ideal. Much of this divergence can be captured by a model that weights sensory information, according only to internal noise in using the cue. These results emphasise the limits of Bayesian models in complex tasks.

---

Imagine that you are trying to give your cat a bath, but as soon as it sees the bathtub, it gets scared and runs away to the garden (Kording et al., 2007; Vilares & Kording, 2011). So, you are walking around your garden, trying to figure out where your cat has hidden, and you hear a “meow” *(auditory cue)*. This perceptual cue is useful but not perfectly reliable and will not allow you to pinpoint exactly the cat’s position. However, from previous experience, you may have learnt that your cat often hides in the bushes, furthest from the pond *(priors)*. The uncertainty in the two pieces of information that you have (the auditory cue and the prior information) allow them to be expressed as probability distributions over location and the optimal strategy for estimating the cat’s location is to integrate the sensory and prior information according to the rules of Bayesian Decision Theory (BDT). Recent studies show that people behave as if they deal with uncertainty in this way, for example, when estimating the position of a hidden target (Berniker, Voss, & Kording, 2010; K.P. Körding & Wolpert, 2004; Tassinari, Hudson, & Landy, 2006; Vilares, Howard, Fernandes, Gottfried, & Kording, 2012), direction of motion (Chalk, Seitz, & Series, 2010), speed (Stocker & Simoncelli, 2006; Weiss, Simoncelli, & Adelson, 2002), or the duration of a time interval (Acerbi, Wolpert, & Vijayakumar, 2012; Ahrens & Sahani, 2011; Jazayeri & Shadlen, 2010; Miyazaki, Nozaki, & Nakajima, 2012). In all of these studies, human observers integrated knowledge of the statistical structure of the experiment (acquired from feedback in previous trials) with sensory information, taking a weighted average according to their relative reliabilities in order to maximise his or her score on the task (Ma, 2012). However, other studies report sub-optimal behaviour, finding that even though observers take into account the uncertainty of the current and prior information, the weights do not match those of an ideal Bayesian observer (Bowers & Davis, 2012; Jones & Love, 2011; Rahnev & Denison, 2018). The fact that human performance ranges from close-to-optimal to largely suboptimal suggests that Bayesian models may describe behaviour well in some cases, but not in others. Understanding when BDT can and cannot provide an accurate model of human behaviour is an important step towards understanding the computations and approximations used by the brain to support adaptive behaviour.

One factor that may influence how close performance is to BDT (“optimal”) predictions is task complexity. For example, Bejjanki, Knill, and Aslin (2016) asked observers to estimate the position of an unknown target (a bucket at a fairground), whose true location was randomly drawn from one of two Gaussian distributions, with different means and variances (*priors)*. On each trial, eight dots were drawn from a Gaussian distribution centered on the true location with either a low, medium or high variance to form a dot cloud that served as a noisy cue to target location (*likelihoods)*. To successfully estimate the position of the target, subjects could use both the likelihood, obtained from the displayed dots, and the prior, obtained from the distribution of previous target positions that they could learn from the trial-by-trial feedback. The study found that subjects adjusted their responses according to the reliability of sensory and prior information, giving more weight to the centroid of the dot cloud (likelihood) when the variance of the prior was high and the variance of the likelihood was low; a signature of probabilistic inference (Ma, 2012). More generally, these results are also in agreement with previously described work, which used *a single prior change* (e.g., Berniker et al., 2010; Vilares et al., 2012). However, unlike in the studies using only a single prior change, the weight placed on the likelihood differed from that of the ideal-Bayes observer whenever the likelihood uncertainty was medium or high. The magnitude of the difference varied with both likelihood and prior uncertainty. As the likelihood became more uncertain, the difference from optimal increased, participants placing more weight on the likelihood than optimal. In addition, the difference from optimal was greater in the narrow prior condition. In a follow-up experiment, participants experienced one prior distribution only, with double the amount of trials used in the original study, finding that subjects’ weights on the likelihoods approach optimal with increasing task exposure, suggesting more time is required to accurately learn the variances of the prior distributions and that learning is disrupted when trying to learn two distributions simultaneously (Bejjanki et al., 2016).

Even in cases when likelihood-weighting might match the prediction of an ideal-Bayes observer, Maloney and Mamassian (2009) noted that such “optimal” or “near-optimal” performance alone is not enough to show that the brain is following Bayesian principles. Maloney and Mamassian (2009) showed that a reinforcement-learning model that learns a mapping from stimulus to response (learning a separate look-up table for each prior and likelihood pairing in the types of tasks discussed here) will also appear to optimally weight prior and likelihood information without learning the individual probability distributions.

Maloney and Mamassian (2009) suggested that the two models may be distinguished by asking whether subjects are able to immediately transfer probabilistic information from one condition to another (hereafter known as *Bayesian transfer*). These transfer criteria are a strong test for use of Bayesian principles because they make very different predictions for how the observer will behave when presented with a new level of sensory noise halfway through the task. If people follow Bayesian principles, we would expect them to *immediately* adapt to the new sensory uncertainty, and integrate it with an already-learnt prior, without any need for feedback-driven learning. On the other hand, the reinforcement-learner would require a certain amount of exposure to the new likelihood and prior pairing (with feedback) in order to form a look-up table that could then lead to optimal performance.

To our knowledge, only one study has tested Bayesian transfer in the context of sensorimotor learning. Sato and Kording (2014) tested the ability of participants to generalise a newly learnt prior to a previously learnt likelihood. In their task, Sato and Kording (2014) first trained participants to complete the task when only a single Gaussian prior was present (either narrow or wide) that could be paired with either a low or high uncertainty likelihood by giving feedback on every trial. After 400 trials, the prior switched to the other level of uncertainty (narrow to wide or wide to narrow) and for the following 200 trials, participants saw the new prior paired only with one of the likelihoods (either low or high) and continued to receive feedback. In the second part of the experiment, subjects still saw the second prior variability, but now with the first likelihood again, which they had so far only seen paired with the first prior. They did not provide any feedback in this part of the task to examine how subjects transferred their knowledge of the prior to the new likelihood. The weight placed on the likelihood in the newly-reintroduced likelihood condition was immediately different to the weight placed on the same likelihood before the change in the prior. In other words, participants’ behaviour in this likelihood condition changed dependent upon prior uncertainty without any explicit training with this prior and likelihood pairing. This is evidence of Bayesian transfer and hence, that participants solved the task using Bayesian principles – representing probabilities – rather than a simpler strategy such as a look-up table learned by reinforcement.

Whether the same will hold in more complex scenarios is unclear. Indeed, it has been repeatedly pointed out that exact Bayesian computations demand considerable computational resources (e.g., working memory, attention) such that the brain might not be able to perform these computations in more complex tasks and will instead resort to heuristics (Beck, Ma, Pitkow, Latham, & Pouget, 2012; Gardner, 2019). We have already seen that performance in more complex tasks is far from optimal (e.g., Bejjanki et al., 2016), suggesting that there are limits to humans’ ability to learn and optimally integrate prior distributions with sensory information when tasks become more complex. Establishing the limits of BDT as a model of human behaviour will inform models of information processing in the human brain.

Here we ask whether people will show Bayesian transfer in a complex situation with multiple priors and likelihood variances, similar to Bejjanki et al. (2016). We report three experiments in which a target is sampled from one of two possible prior distributions (with different means and different variances) and then cued with one of three possible likelihood variances (with the variance itself also displayed). Likelihood and prior variances were identical to those used in Bejjanki et al. (2016) in terms of visual angle, in order to match the true (objective) reliabilities of the cue and prior across the studies. The first two experiments tested for Bayesian transfer by only introducing the high likelihood variance in blocks 4 and 5 of the task. The only difference was that in the second experiment, participants were explicitly told that the prior variances differed, to test whether this would promote closer-to-optimal performance. The last experiment was used to check whether removing the additional burden of transfer allows participants to learn the complex environment correctly by presenting all likelihood conditions from the start of the experiment – a replication of Bejjanki et al. (2016).

To summarise, in the first experiment, we found that observers did not show evidence for Bayesian transfer. When a new high variance likelihood was introduced in blocks 4 and 5, they did not weight it less than the familiar medium variance likelihood. This is at odds with the idea that observers perform full Bayesian inference, combining prior and likelihood based on relative variances, and thus does not provide evidence for Bayesian transfer. In the second experiment, observers did show evidence for transfer, weighting the newly-introduced high variance likelihood significantly less than the medium variance likelihood. However, in both experiments, the weights placed on the medium and high variance likelihoods were much higher than optimal. These weights remained sub-optimal in the final experiment where all likelihood variances were present from the start of the task. These results extend our knowledge of how potentially-Bayesian perceptual processes function in complex environments.

## Experiment 1: Testing transfer to a new level of likelihood variance

In the first experiment, we tested whether Bayesian transfer would occur in a complex environment with two priors, similar to the one used by Bejjanki et al. (2016). We trained participants on a spatial localization task with two likelihood variances and two prior distributions (with different means and variances). In initial training, they were exposed to all four combinations (trials interleaved), with feedback. If, like participants in Bejjanki et al. (2016), they weighted the likelihood and the prior differently across conditions in line with their differing reliabilities, this would show that they had learned and were using the priors. However, such reliability-weighting could either be done via Bayesian inference – representing probabilities – or via a simpler strategy akin to learning a look-up table (Maloney & Mamassian, 2009). To distinguish these possibilities, after the training trials, we tested for “Bayesian transfer” by adding a new higher-variance likelihood distribution to the task. If participants deal with this newly-introduced likelihood in a Bayesian manner, they should immediately rely less on this new likelihood information than they did on the likelihoods in previously-trained conditions. Alternatively, if their initial learning is more rote in nature (i.e., more like a look-up table), participants would begin to place a different weight on the new likelihood only after extensive training with feedback.

### Methods

#### Overview

Subjects performed a sensorimotor decision-making task on a computer monitor where they estimated the horizontal location of a hidden octopus. The true location was sampled from one of two distinct Gaussian distributions that differed in mean and variance (narrow or wide priors). On each trial, the relevant prior distribution was indicated by the colour of the likelihood information – eight dots that were described to the participant as the “tentacles” of the octopus. The horizontal locations of the eight dots were sampled from a Gaussian distribution centred on the true location that had either low, medium, or high variability (the likelihood). To estimate the octopus’ position, participants could use (although this was never explicitly mentioned) both the likelihood and prior information, with the subjects able to learn the latter via trial-to-trial feedback. Participants completed five blocks of trials. Crucially, in blocks one to three only the low and medium likelihood variances were paired with the narrow or wide priors. The high likelihood condition was only introduced in blocks four and five to test for evidence of Bayesian transfer.

#### Participants

Participants were recruited from the Durham Psychology department participant pool, Durham University newsletter, and by word-of-mouth. Twenty-six participants were recruited in total (13 female, mean age: 20.1, age range: 18-30 years). All participants had normal or corrected-to-normal visual acuity, and no history of neurological or developmental disorders. Each participant received either course credits or a cash payment of £10 for their time.

#### Ethics

Ethical approval was received from the Durham University Psychology Department Ethics Board. All participants gave written, informed consent prior to taking part in the study.

#### Stimuli and apparatus

Stimuli were displayed on a 22-inch iiyama monitor (1680 × 1050 pixels), viewed at a distance of 60cm, using the Psychophysics Toolbox for MATLAB (Brainard, 1997; Kleiner et al., 2007; Pelli, 1997). The stimuli were set against a blue background (to represent the sea).

The position of the octopus was sampled from one of two Gaussian distributions (priors): the narrow (standard deviation of *σ*_*pl*_ = 1% of screen width) or wide (standard deviation of *σ*_*ph*_ = 2.5% of screen width) priors. The side of the screen associated with each prior was counterbalanced across participants. One was always 35% of the way across the screen (from left to right), and the other 70%. When the narrow prior was centered on 35%, for example, the wide prior had a mean in the opposite side of the screen (i.e., to the right, centered on 70%). When the octopus appeared on the left-hand side (drawn from the prior centered on 35%) it was white, and when it appeared on the right (drawn from the prior centered on 70%) it was black.

At the beginning of each trial, a cloud of eight dots (0.5% of screen width in diameter) appeared on the screen. The horizontal position of each dot was drawn from a Gaussian distribution centered on the true octopus location with either low (*σ*_*ll*_ = 0.6% screen width), medium (*σ*_*lm*_ = 3% screen width), or high (*σ*_*lh*_ = 6% screen width) standard deviation (in the following referring to as low, medium and high variance likelihood conditions). The horizontal positions of the dots were scaled so that their standard deviation (SD) was equal to the true SD (*σ*_*ll*_, *σ*_*lm*_ or *σ*_*lh*_) on each trial while preserving the mean of the dots. We performed this correction so that participants would “see” the same variability across trials for each likelihood condition. This ensures that an observer who computes the reliability for the likelihood information trial by trial would always calculate the same value within likelihood trial types. The vertical positions of the dots were spaced at equal intervals from the vertical center of the screen, with half of the dots appearing above, and the other half, below the center. The vertical distance between each dot was fixed and equal to 1% screen width). Given that the vertical positions of the dots were fixed, only the horizontal position of the target was relevant. Participants estimated location only along the horizontal axis by moving a vertical green rectangle (measuring 1% of screen width in width and 3% of screen width in height) left or right, making this a one-dimensional estimation task. Participants received feedback in the form of a red dot (0.5% of screen width in diameter) that represented true target position.

The combination of two priors and three likelihoods led to six trial types (all possible prior × likelihood pairings). The task was split into five blocks of trials with 300 trials per block. In the first three blocks of the task only four trial types were used (75 trials of each paring), with the high likelihood condition not shown in combination with either prior. The high likelihood condition was introduced in blocks four and five (50 trials per paring), in order to test for Bayesian transfer. Within each block, all trial types were interleaved. The trials were broken into runs of 20 trials. Within each run the trials for each prior type were arranged such that an ideal learner would have an exact estimate of the mean and variance of the prior distributions if evidence was accumulated over those 20 trials.

We also included prior-only trials where subjects were told that a black/white octopus was hiding somewhere on the screen and instructed to find it. No sensory information was provided (no likelihood information). These trials were interleaved with the rest of the trials (one every 9 trials for each prior), and there were 83 trials in total, for each of the priors (narrow and wide).

#### Procedure

Participants were instructed to estimate the position of a “hidden” octopus, indicating their estimate by adjusting the horizontal location of a “net” (green rectangle). Each trial started with the presentation of eight dots that remained on screen until the end of the trial (the likelihood information, described to the participants as the tentacles of the octopus) (Figure 1A). The eight dots could have one of three levels of uncertainty: low, medium, or high variance likelihood trials (Figure 1B). When the level of uncertainty was higher, the dots were more dispersed on the screen and, therefore, were a less reliable indicator of the true location of the octopus. Participants used the mouse to move the net to their guessed position, using a right click to confirm their choice (no time limit). Following a response, the true position of the octopus was shown as a red dot on the screen. Over the course of the experiment, the feedback served as a second cue to location since the true locations of the black and white octopi were drawn from different distributions. In other words, participants could learn a prior over each octopus’ location. We provided performance feedback on a trial-to-trial basis so that the priors could be learned. Specifically, subjects could potentially learn that the two sets of octopi (black/white) were drawn from separate Gaussian distributions centred at different locations on the screen and with differing levels of uncertainty (narrow and wide variance prior trials).

**Figure 1.**
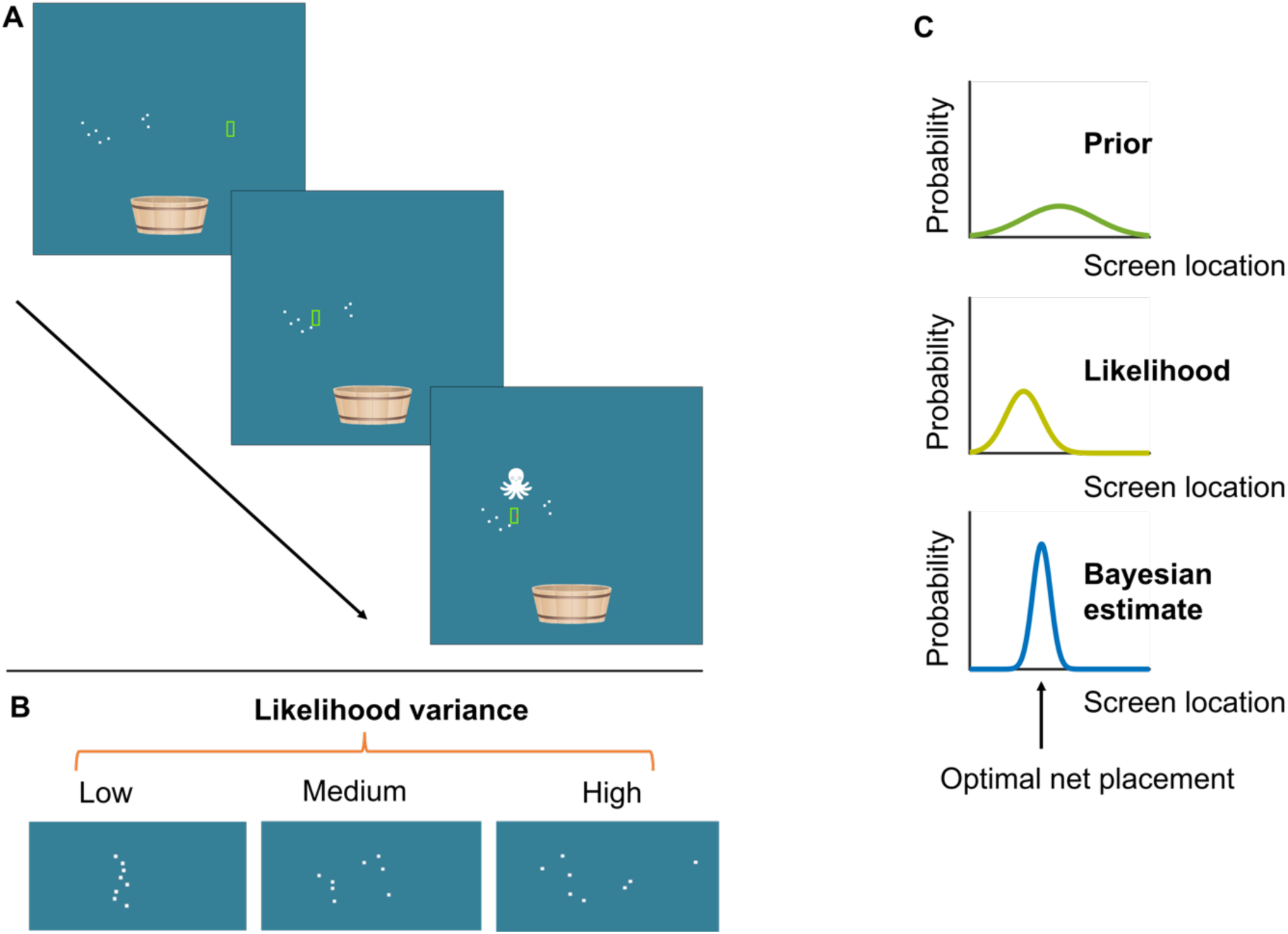
A) Illustration of the task. Participants were asked to estimate the position of a hidden target (the “octopus”, represented as the red dot) by horizontally moving a net (green vertical bar). At the beginning of each trial, participants were given noisy information about the location of the hidden target in the form of eight dots (the likelihood). Participants then moved the net to the estimated location and clicked to confirm their response, after which the actual target location was displayed. If the target was inside, or overlapped with, the net, a score was added to the participant’s score. B) The three likelihood variances. C) Illustration of a Bayes-optimal observer. A Bayesian observer would combine information about the prior uncertainty (learnt from the distribution of previous target locations) with the likelihood information on a given trial to optimally estimate the target location.

To keep participants engaged, we incorporated an animation when the participant picked the right location of a cartoon octopus moving into a bucket centred at the bottom of the screen. In addition, participants would get one point added to their score if they “caught” the octopus. An octopus was caught if the true octopus position overlapped with the net placement by at least 50% of the red feedback dot’s size. The cumulative score was displayed at the end of each trial. Participants completed 5 blocks of 300 trials each for a total of 1500 trials. The five experimental blocks were performed in succession with a short break between each one. The experiment duration was approximately an hour and a half.

#### Data Analysis

For each individual participant, we regressed estimated octopus position against the centroid (mean) of the cloud of dots (likelihood) on each trial. All regression analyses were done using a least squares procedure (the polyfit function in MATLAB). The slope of the fitted regression line quantifies the extent to which participants rely on the current sensory evidence (likelihood), as opposed to prior information. A slope of one suggests that participants only use likelihood information and a slope of zero suggests that participants rely only on their prior knowledge, ignoring the likelihood. A slope between zero and one suggests that both likelihood and prior information are taken into account, and the steeper the slope, the more participants rely on the likelihood and less on the prior information. Accordingly, we will refer to the fitted slope values as the weight placed on the likelihood.

We also computed the weight that would be given to the likelihood in each condition by an ideal Bayesian observer with perfect knowledge of the prior and likelihood distributions (see Figure 1C for an illustration). The optimal weight on the likelihood was computed as:

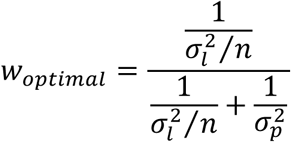

where 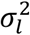 is the variance of the likelihood, *n* is the number of dots that indicate the likelihood (in this case, there were 8 dots), and 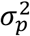 is the variance of the prior.

To determine the proportion of the variance in responses that is accounted for by change in the estimate from the sensory cue, the coefficient of determination (*R*^2^) was calculated by linearly regressing participants’ responses against each estimate participants could have taken from the cue (i.e., arithmetic mean, robust average, median or mid-range). This was done for the combined data of all subjects in each experiment, across all blocks and trial types (prior and likelihood pairings). The estimate with the highest *R*^2^ value was taken to be the estimate participants had most likely used.

Statistical differences were analysed using repeated-measures ANOVA with a Greenhouse-Geisser correction (Greenhouse & Geisser, 1959) of the degrees of freedom in order to correct for violations of the sphericity assumption if ϵ ≤ 0.75 and a Huynh-Feldt correction otherwise.

We discarded a trial from analysis if the absolute error for that trial was in the top 1% of all absolute errors, computed separately for each prior and likelihood pairing across all blocks and participants (this rule excluded at most 13 trials per pairing for an individual subject).

#### Results and Discussion for Experiment 1

We first checked whether subjects took the mean as an estimate from the sensory cue, and not a heuristic, such as the robust average. In tasks similar to ours (Bejjanki et al., 2016; Chambers, Gaebler-spira, & Kording, 2018; Vilares et al., 2012), authors assume that observers use the mean of the dots as their best estimate of true location from the likelihood information. However, we did not explicitly tell our participants how the eight dots that formed the likelihood were generated, or that the best estimate they could take from them was their mean position, leaving open the possibility that observes may have taken a different estimate from the cue than the mean (de Gardelle & Summerfield, 2011; Van Den Berg & Ma, 2012). The mean horizontal position was found to explain the most amount of variance in participants’ responses (*R*^2^ = 0.996), relative to the robust average (*R*^2^ = 0.995), median (*R*^2^ = 0.995) or the mid-range of the dots (*R*^2^ = 0.992). This suggests that the mean of the dots is the estimate that participants take from the sensory cue.

We then examined whether the weight participants placed on the likelihood, relative to the prior, varied with respect to trial type (prior/likelihood pairing) for all the trial types present from the beginning of the experiment. Without this basic result – a replication of the pattern found by Bejjanki et al. (2016) – we could not expect them to transfer knowledge of the learnt prior distributions to the new high variance likelihood in the later blocks. This was a qualified success: for these trial types (blue and green bars in Figure 2), participants showed the predicted pattern, but placed more weight on the likelihood than is optimal (compare bar heights to dashed lines in Figure 2), in line with previous research (Bejjanki et al., 2016; Tassinari et al., 2006). We conducted a 2 (narrow versus wide variance prior) × 2 (low versus medium variance likelihood) × 5 (block) repeated measures ANOVA with the weight given to the likelihood (the displayed dots) as the dependent variable. These results are shown in Table 1 and summarised here. There was a main effect of prior variance, with less weight on the likelihood when the prior was narrower (*p* < .001). There was also a main effect of likelihood variance (*p* < .001), where participants relied less on the medium variance likelihood. However, there was also a significant interaction effect of likelihood and prior (*p* = .001). When the prior was narrow, the decrease in reliance on the likelihood was smaller as the likelihood variance increased (*t*(25) = 3.57, *p* = .001).

**Table 1.** Results from a 2 (prior) × 2 (likelihood) × 5 (block) Repeated Measures ANOVA for the likelihood variances present from the beginning of the task (low and medium) in Experiment 1

| | *F* | *df, df<sub>error</sub>* | *p* | Effect size ($\eta_p^2$) |
| --- | --- | --- | --- | --- |
| Likelihood | 104.40 | 1, 25 | <.001 | .81 |
| Prior | 28.08 | 1, 25 | <.001 | .53 |
| Block | 7.72 | 4, 100 | <.001 | .24 |
| Likelihood x prior | 12.77 | 1, 25 | .001 | .34 |
| Likelihood x block | 3.32 | 4, 100 | .01 | .12 |
| Prior x block | .51 | 4, 100 | .21 | .06 |

**Figure 2.**
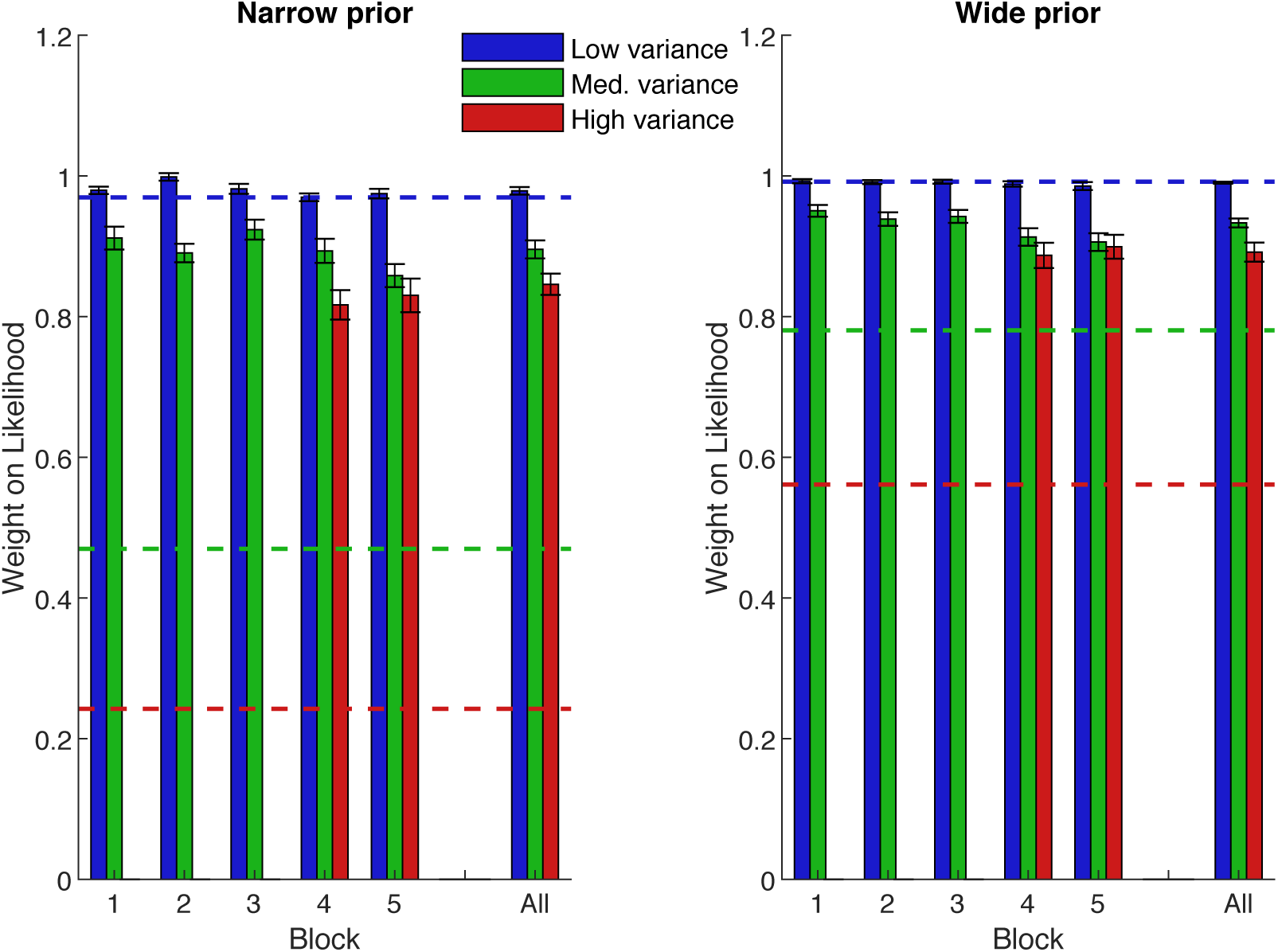
Mean weight placed on the likelihood information, separated by block and prior width in Experiment 1. Lower values represent a greater weight on the prior. Blue is low-variance likelihood (a tight array of dots), green is medium-variance likelihood (dots somewhat spread out), red is the later-introduced high-variance likelihood (highly spread out dots). Dashed lines show optimal-predicted values. Error bars are +/− 1 SEM. The far right is the average over blocks.

We found a main effect of block (*p* < .001) and an interaction between block and likelihood (*p* = .014), with the medium variance likelihood weighted significantly differently across blocks (simple main effect of block, *F*(4,100) = 5.84, *p* < .001, 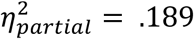, weights decrease with increasing exposure), but not the low variance likelihood (no simple main effect of block, *F*(4,100) = 1.64, *p* = .169, 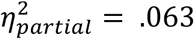). This suggests that participants adjusted, through practice, their weights on the medium variance likelihood, getting closer to optimal.

Examination of the prior-only trials shows successful learning of the priors. On average, subjects’ responses were not significantly different from the prior mean in the wide prior condition (*t*(25) = −0.77, *p* = .450). They were significantly different in the narrow prior condition (*t*(25) = −2.78, *p* = .010), although the bias was extremely small (95% CI: [0.06,0.41] percent of the screen width to the left). The median standard deviation of responses for all subjects was 1.4% (narrow prior) and 2.5% (wide prior): almost identical to the true prior SDs of 1.3% and 2.5%, respectively.

Participants qualitatively followed the predicted optimal pattern of reweighting: like the dashed lines (predictions) in Figure 2, actual likelihood weights (bars) were higher for the wide prior (right) than the narrow prior (left), and higher for the low variance likelihood (blue) than the medium variance likelihood (green). However, comparing bar heights with dashed lines (predictions) shows that quantitatively, their weights were far from optimal. Participants systematically gave much more weight than is optimal to the likelihood when its variance was medium (Figure 2, green bars vs lines – *p* < .001 in all blocks for the medium likelihood when paired with either prior). This over-reliance on the likelihood is in line with previous studies (e.g., Bejjanki et al., 2016), although stronger in the present study. Participants, therefore, accounted for changes in the probabilities involved in the task (e.g., weighted the likelihood less when it was more variable), but did not perform exactly as predicted by the optimal strategy.

Having found that participants’ performance was in line with the predicted patterns, we could then ask if they would generalise their knowledge to the new high likelihood trials added in blocks 4-5 (“Bayesian transfer”), as predicted for an observer following Bayesian principles. This should lead immediately to a lower weight for the new high variance likelihood than the familiar medium variance likelihood. By contrast, lack of a significant difference in weights between the medium and high likelihood trial types would suggest that the observer is employing an alternative strategy, such as simply learning a look-up table. To test this, we performed a 2 (prior) × 3 (likelihood) × 2 (block) repeated measures ANOVA (summarise only blocks 4 and 5 – those with all likelihoods present). These results are shown in Table 2 and summarised here. As above, we found a main effect of likelihood, participants placing less weight on the likelihood as it became more uncertain (*p* < .001). However, post-hoc analyses showed that the weight placed on the high likelihood was not significantly lower than the weight placed on the medium likelihood (*p* = .103). Only the comparison of the weights placed on the likelihood in low and high variance trial types was significant (*p* < .001). Moreover, there was no main effect of block (*p* < .28), nor an interaction between block and likelihood (*p* = .48), suggesting that the weight placed on the newly introduced likelihood variance did not decrease with increasing exposure.

**Table 2.** Results from a 2 (prior) × 3 (likelihood) × 2 (block) Repeated Measures ANOVA for all likelihood variances in Experiment 1

| | $F$ | $df, df_{error}$ | $p$ | Effect size ($\eta_p^2$) |
| --- | --- | --- | --- | --- |
| Likelihood | 50.08 | 2, 50 | <.001 | .67 |
| Prior | 15.52 | 1, 25 | .001 | .38 |
| Block | 1.21 | 1, 25 | .28 | .05 |
| Likelihood x prior | 2.39 | 1.62, 40.41 | .10 | .09 |
| Likelihood x block | .75 | 2, 50 | .48 | .03 |
| Prior x block | 4.35 | 1, 25 | .05 | .15 |

Finally, we compared mean weights in block 5 against the optimal Bayesian values for each prior and likelihood pairing. In the low variance likelihood trials we did not observe significant deviation from the Bayesian prediction, irrespective of prior (low variance likelihood, narrow prior: *t*(25) = .784, *p* = .440); low variance likelihood, wide prior: *t*(25) = −1.12, *p* = .270). Subjects’ weights differed significantly from optimal in all other conditions (*p* < .001 in all cases).

Overall, our results do not exactly match the predictions of a Bayesian observer because we find only weak evidence of Bayesian transfer. Specifically, while we find a main effect of likelihood, the weight on the high variance likelihood is not significantly different to that placed on the medium variance likelihood (although the change is in the predicted direction). That said, our results are not simply more consistent with a rote process, since the weight placed on the high likelihood does not decrease with increasing exposure (no interaction between likelihood and block).

Our results point mostly away from a simple variance weighted Bayesian model being a good model of human behaviour in this particular task. The correct pattern of weights was present, but evidence of transfer was weak. Participants were also significantly sub-optimal, overweighting the likelihood whenever its variance was medium or high. Previous studies have also found that observers give more weight to the sensory cue than is optimal (e.g., Bejjanki et al., 2016); even so, the level of sub-optimality that we observe here is still drastically higher, compared to previous reports. However, Sato and Kording (2014) found better, near-optimal performance in those participants who were told that the sensory information can have one of two levels of variance, and that the variance will sometimes change, compared to those who were not provided with this information. We therefore reasoned that if observers are given additional information about the structure and statistics of the task (e.g., that the variances of the prior distributions are different), the weight they give to the sensory cue may move closer to optimal. If we find weights closer to optimal, we may be better able to detect whether transfer had taken place because the effect size of a change in the likelihood would be bigger. In fact, we wonder whether the size of this effect could be an important factor behind the lack of significant differences observed in Experiment 1, i.e., that the effect size of the change from medium to high likelihood was too small for our statistical analysis to reliably detect. In view of this, we set out to test whether additional instructions will lead to weighting of likelihood and prior information that is closer to optimal.

## Experiment 2: Additional instructions about prior variance

Experiment 2 was identical to Experiment 1 except for a change in instructions. In this experiment, subjects were explicitly (albeit indirectly) informed of the different variances of the prior. We hypothesised that giving participants additional information about the model structure of the task will move weights closer to optimal and make any transfer effects more pronounced.

### Methods

Twelve participants (8 female, mean age: 20.3, age range: 19-22 years) participated in Experiment 2. All participants had normal or corrected-to-normal visual acuity, no history of neurological or developmental disorders and had not taken part in Experiment 1. Each participant received either course credits or cash compensation for their time.

The experimental set had the same layout as the main experiment, with the following difference: in addition to the previously described instructions, subjects in this version of the task were told that “it is important to remember that one of the octopuses tends to stay in a particular area, whereas the other one moves quite a bit!” (i.e., they were indirectly informed that the variances of the priors were different). (see Appendix A for full instructions).

### Results and Discussion for Experiment 2

Similarly to what we saw in Experiment 1, the mean and robust average of the dots explained the same amount of the variance in participants’ responses (*R*^2^ = 0.991 for both), followed by the median (*R*^2^ = 0.990) and the mid-range (*R*^2^ = 0.989). We, thus, proceed with the mean as the estimate from the likelihood.

Figure 3 shows that the pattern of results was qualitatively similar to those of Experiment 1 (see Figure 2). A 2 (prior) × 2 (likelihood) × 5 (block) repeated measures ANOVA (analysing only the low and medium likelihood trials) revealed that subjects placed less weight on the likelihood as its uncertainty increased (main effect of likelihood, *p* < .001) and as the prior uncertainty decreased (main effect of prior, *p* = .002). However, unlike in Experiment 1, there was no significant interaction of these factors (*p* = .123) (see Table 3).

**Table 3.** Results from a 2 (prior) × 2 (likelihood) × 5 (block) Repeated Measures ANOVA for the likelihood variances present from the beginning (low and medium) in Experiment 2

| | $F$ | $df, df_{error}$ | $p$ | Effect size ($\eta_p^2$) |
| --- | --- | --- | --- | --- |
| Likelihood | 46.14 | 1, 11 | <.001 | .81 |
| Prior | 17.28 | 1, 11 | .002 | .61 |
| Block | 3.15 | 4, 44 | .02 | .22 |
| Likelihood x prior | 2.80 | 1, 11 | .12 | .20 |
| Likelihood x block | 3.73 | 4, 44 | .01 | .25 |
| Prior x block | 1.32 | 2.07, 22.77 | .29 | .11 |

**Figure 3.**
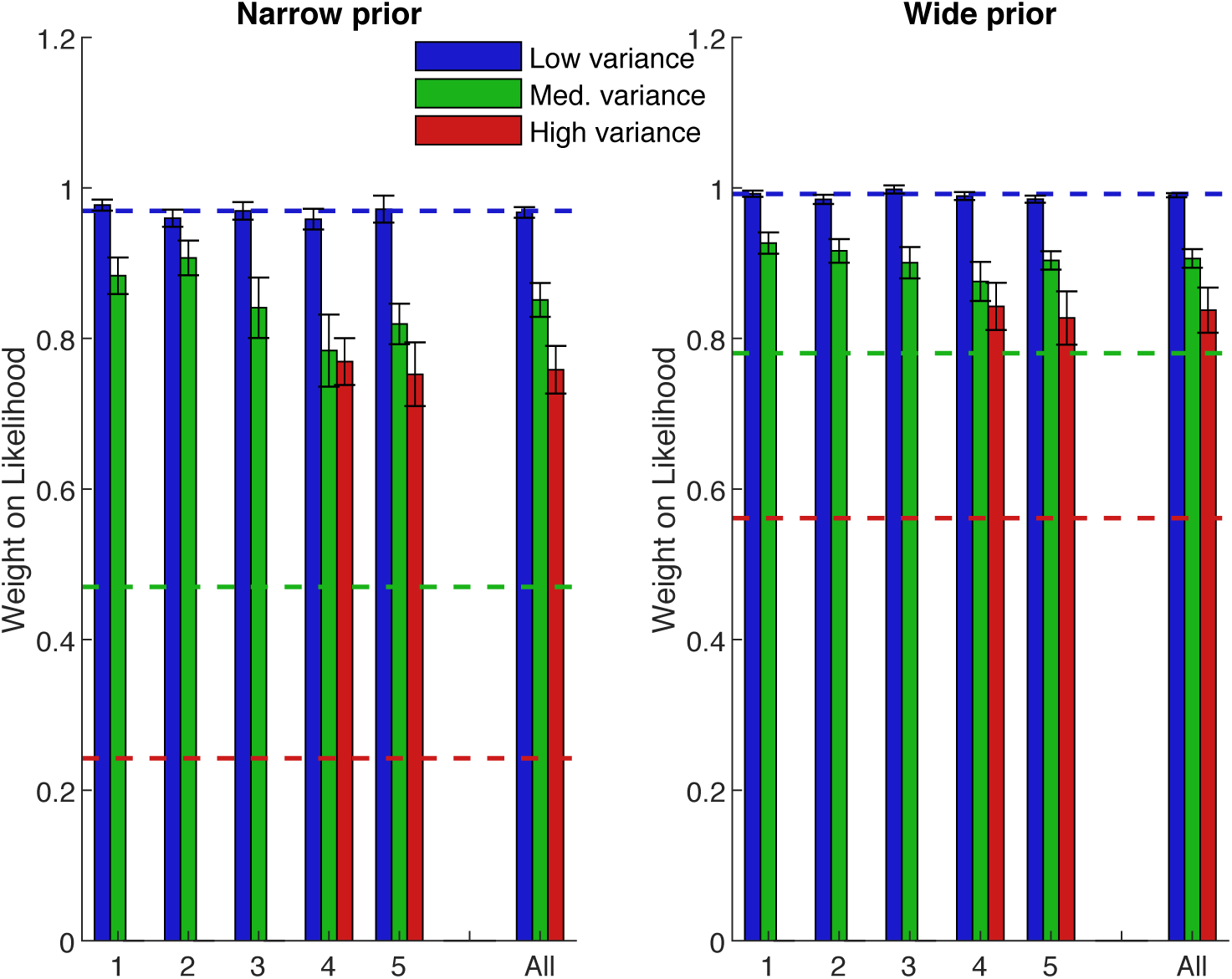
Mean weight placed on the likelihood information in each block of Experiment 2. Blue is the low-variance likelihood, green is the medium-variance likelihood, red is the high-variance likelihood. Dashed lines show optimal values. Error bars are +/− 1 SEM. The far right is the average over blocks.

We found a main effect of block (*p* = .02) and an interaction between block and likelihood (*p* = .01), with participants weighting the likelihood significantly less with increasing task exposure (regardless of prior) when its variance was medium (*F*(2.21,25.34) = 3.81, *p* = .03, 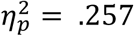, with a Greenhouse-Geisser correction), but not when it was low (*F*(4,44) = 0.70, *p* = .60, 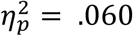).

As before, we analysed subjects’ responses in the prior-only trials, finding a good quantitative agreement with the true prior mean (narrow prior: *t*(11) = −0.002, *p* = .999; wide prior: *t*(11) = 0.35, *p* = .734). The median standard deviation (SD) of responses was also remarkably similar to the true prior SDs (narrow prior: 1.6% vs. 1.3% in screen units; wide prior: 2.5% for both).

Again, subjects’ overall performance was suboptimal (as can be seen by comparing the height of the bars against the dashed lines – the optimal predictions – in Figure 3). Subjects’ placed more weight on both the medium and high variance likelihoods than is optimal (*p* < .001 in both cases, for both priors). However, it is worth noting that the weights placed on the medium and high likelihoods are closer to optimal than they were in Experiment 1 (compare bar heights in Figure 2 and 3).

Finally, we tested for transfer to the newly-introduced high likelihood in blocks 4-5. We conducted a 2 (prior) × 3 (likelihood) × 2 (block) repeated measures ANOVA (analysing only blocks 4 and 5 with all likelihoods present). These results are shown in Table 4 and summarised here. There was a main effect of likelihood, with less weight placed on the likelihood as it became more uncertain (*p* < .001). Unlike in Experiment 1, post-hoc analysis showed that the weight placed on the high likelihood was significantly lower than the weight placed on the medium likelihood (*p* = .034). The weights placed on the likelihood in the low variance trial type were significantly lower than those in the medium and high variance trial types (*p* < .001 for both). Moreover, there was no main effect of block (*p* < .64), or an interaction effect of block and likelihood (*p* < .15), meaning that the weight placed on the newly-added likelihood information did not vary with increasing exposure.

**Table 4.** Results from a 2 (prior) × 3 (likelihood) × 2 (block) Repeated Measures ANOVA for all likelihood variances in Experiment 2

| | $F$ | $df, df_{error}$ | $p$ | Effect size ($\eta_p^2$) |
| --- | --- | --- | --- | --- |
| Likelihood | 37.98 | 2, 22 | <.001 | .78 |
| Prior | 18.36 | 1, 11 | .001 | .63 |
| Block | .23 | 1, 11 | .64 | .02 |
| Likelihood x prior | 1.80 | 2, 22 | .19 | .14 |
| Likelihood x block | 2.09 | 2, 22 | .15 | .16 |
| Prior x block | .17 | 1, 11 | .69 | .02 |

Again, we find a significant difference between subjects’ weights (in block 5) and optimal predictions when the likelihood variance was medium or high (*p* < .001 in both cases), but not when it was low, irrespective of prior variance (low likelihood, narrow prior: *t*(11) = .120, *p* = .907); low likelihood, wide prior: *t*(11) = −1.29, *p* = .163).

In line with our prediction of transfer, here we show that the observers put lower weight on the high variance likelihood than the medium variance likelihood they have experienced before. This is strengthened by the fact that participants’ weights did not change significantly with increasing exposure across blocks 4-5.

To check more directly for differences due to experimental instructions, we compared subjects’ performance in the last 2 blocks across the two experiments. We ran a 2 (instructions) × 2 (prior) × 3 (likelihood) × 2 (block) mixed ANOVA. We found a main effect of instructions (*F*(1,36) = 6.78, *p* = .013, 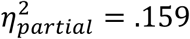) with subjects weighting the likelihood significantly less with explicit instructions (Experiment 2), i.e. closer to the optimal weightings. We also found an interaction between instructions and likelihood (*F*(1,36) = 4.79, *p* = .011, 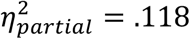), indicating that the main effect of instructions is due to a significant decrease in the weight placed on the high, relative to the medium likelihood in the explicit instructions (Experiment 2) task, but not the original task (Experiment 1).

These results show that adding extra instructions to the task that make the participant aware of a change in uncertainty between the two priors has an effect. The weights placed on the likelihood moved closer to optimal, and the transfer criterion was met, which suggests that, perhaps, observers are more likely to adopt a Bayes-optimal strategy when more explicit expectations about the correct model structure of the task are set. However, even with the additional instructions, the weight given to the sensory cue was still systematically higher than the “optimal” weight. Arguably, expecting people to perform optimally is rather unrealistic, as it presumes that the observer perfectly knows the environmental statistics. However, Bejjanki et al. (2016) found performance much closer to optimal than what we have seen in either of our previous experiments. The major difference between their experiment and ours’ is the fact that Bejjanki et al. (2016) presented all likelihood variances from the start of the task. Therefore, unlike Experiments 1 and 2, which were designed in order to provide some evidence of transfer, Experiment 3 sought to test whether subjects’ weights would move closer to optimal if we present all likelihood variances from the beginning, in a more direct replication of Bejjanki et al. (2016). The likelihood and prior variance parameters were identical to those used in Bejjanki et al. (2016), and we used a similar number of trials per prior and likelihood pairing (250 vs. 200 trials in Bejjanki et al. (2016).

## Experiment 3: All likelihoods from the beginning

Experiment 3 was identical to Experiment 1 (lacking the extra instructions of Experiment 2) except that all likelihood variances were included from the beginning of the task. The participants experienced all six trial types in every block.

### Methods

Twelve participants (10 female, mean age: 22.6, age range: 19-30 years) took part in Experiment 3. All participants had normal or corrected-to-normal visual acuity, no history of neurological or developmental disorders and had not taken part in Experiment 1 and 2. Each participant received either course credits or cash compensation for their time.

The stimuli and task were identical to those described for Experiment 1, except that all likelihood conditions (low, medium and high) were now present from the beginning (50 trials of each likelihood/ prior pairing interleaved in the same block).

### Results and Discussion of Experiment 3

Again, the mean position of the dots explained the most amount of variance in participants’ responses (*R*^2^ = 0.990). The amount of variance explained decreased for the robust average (*R*^2^ = 0.989), median (*R*^2^ = 0.988) an the mid-range of the dots (*R*^2^ = 0.985). We, thus, proceed with the mean as the estimate from the likelihood.

Figure 4 shows a similar pattern of results to Experiments 1 and 2. Again, a 2 (prior) × 3 (likelihood) × 5 (block) repeated measures ANOVA shows that the likelihood information was weighted less as it became more unreliable (main effect of likelihood, *p* < .001). Specifically, subjects placed significantly more weight on the low likelihood than on the medium (*p* = .001) or high likelihood (*p* < .001), and more weight on the medium likelihood than the high likelihood (*p* = .005). No other main effects or interactions were significant (see Table 5 for a summary of results).

**Table 5.**
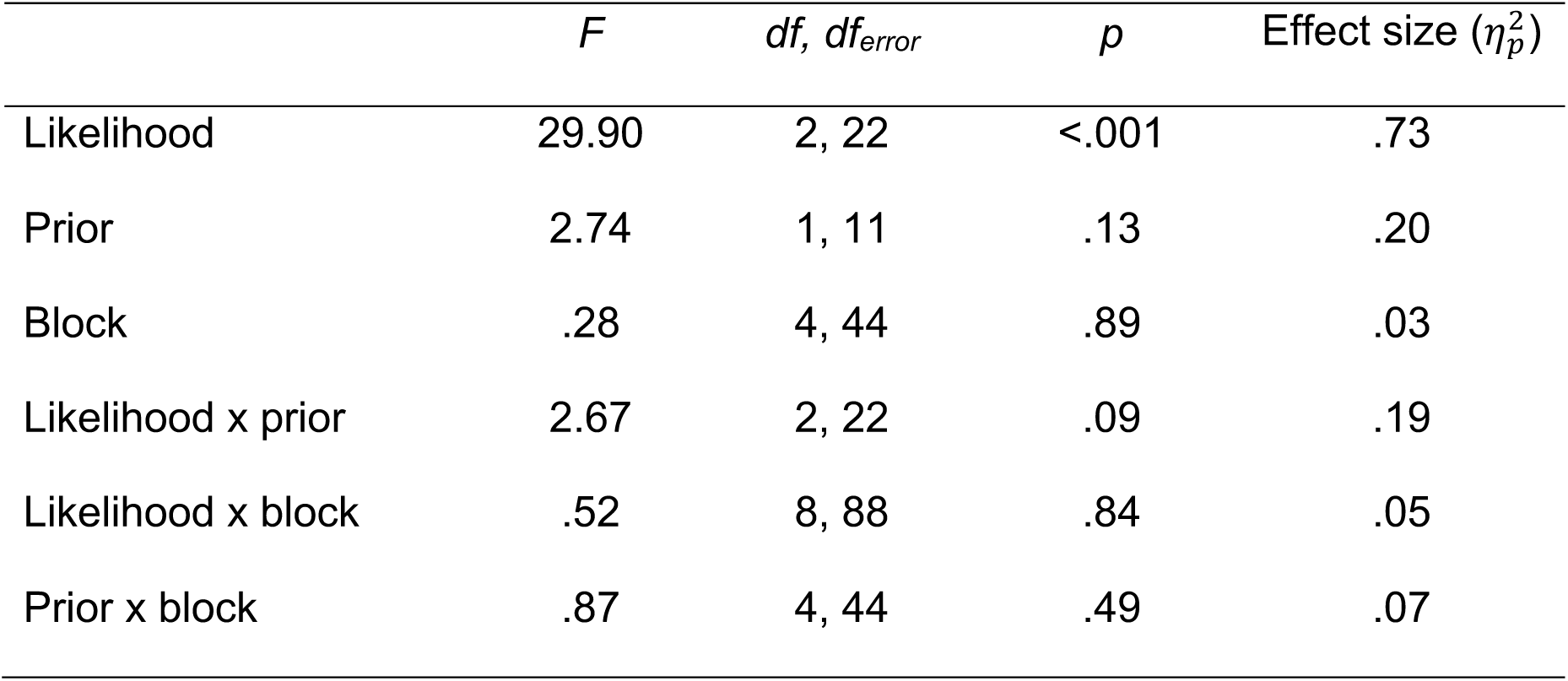
Results from a 2 (prior) × 3 (likelihood) × 5 (block) Repeated Measures ANOVA for all likelihood variances in Experiment 3

**Figure 4.**
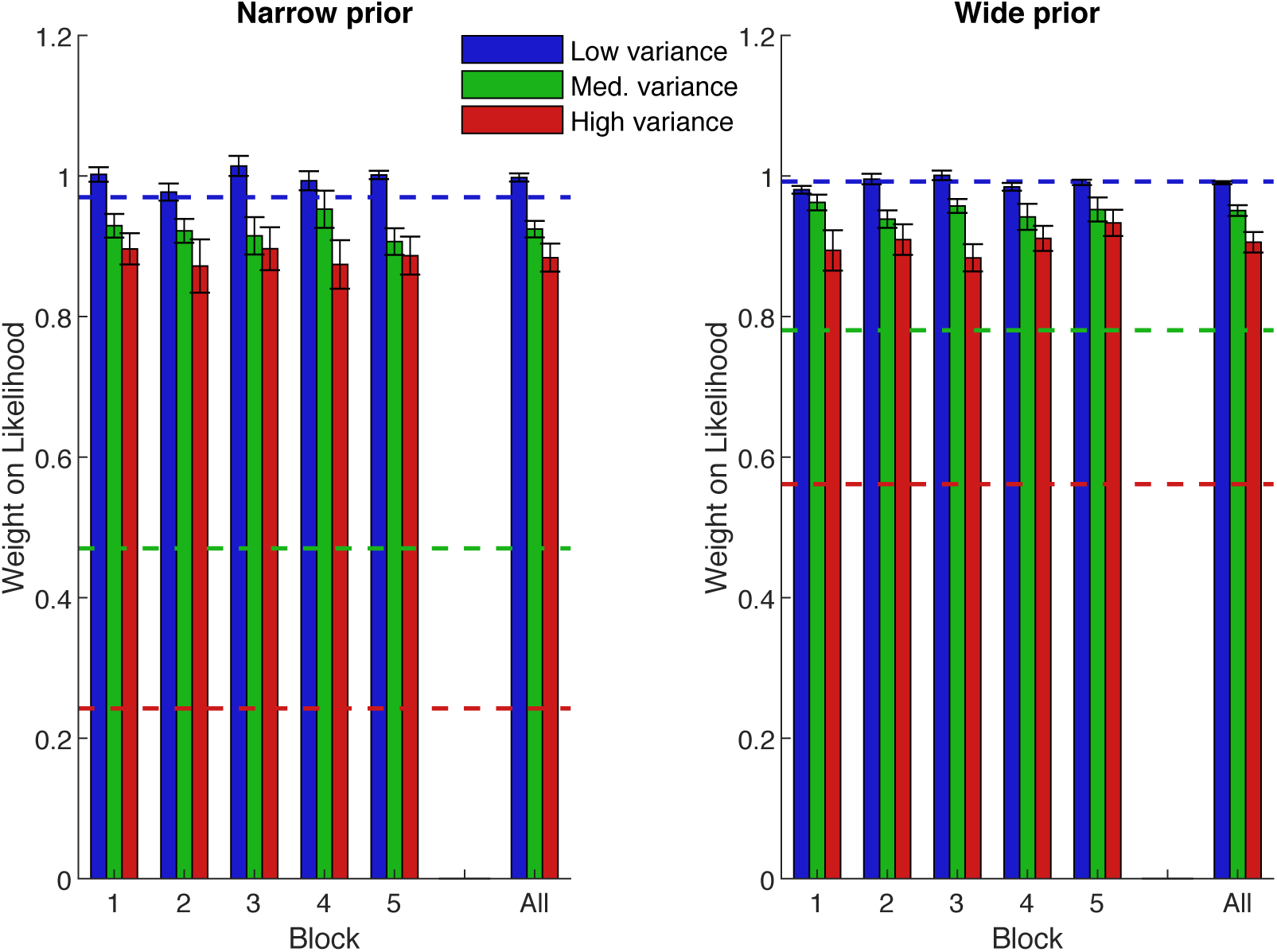
Mean weight placed on the likelihood information in each block of Experiment 3. Blue is the low-variance likelihood, green is the medium-variance likelihood, red is the high-variance likelihood. Dashed lines show optimal values. Error bars are +/− 1 SEM. The far right is the average over blocks.

For the prior-only trials, subjects’ responses were, on average, statistically indistinguishable from the mean of the wide prior distribution (*t*(11) = −1.14, *p* = .278), but were significantly different from the mean of the narrow prior (*t*(11) = −3.91, *p* = .002) (although we note that the bias was small (95% CI: [0.24,0.87] percent of the screen width to the left). The median standard deviation (SD) of responses was 2.2% for the narrow prior condition and 2.6% for the wide prior condition; the SD of responses was, therefore, only close to the true variance of the wide prior (which was 2.5%). Together, these findings suggest that subjects had not learnt either the mean, or the variance of the narrow prior condition. This may explain the lack of difference in performance between the narrow and wide prior conditions in this task.

A comparison of subjects’ weights on the likelihoods in block 5 against Bayesian predictions showed a significant difference for all likelihood and prior pairings (*p* < .001), with the exception of the wide prior/ low likelihood condition (*t*(11) = −.362, *p* = .724).

To sum up, although the correct pattern of weights was present, subjects were still substantially sub-optimal, even after experiencing all likelihood variances from the start of the task.

### Accounting for Suboptimality

Even when we replicate Bejjanki et al. (2016) very closely with all likelihoods from the beginning, our participants are strikingly suboptimal. However, we note that in our initial calculations of optimal behaviour, we assumed that observers weight sensory and prior information according only to the variance of the dot distribution (i.e., external noise). However, many of the studies in the cue combination field that have found near-optimal performance used cues that only have internal noise, not external (Alais & Burr, 2006; Körding & Wolpert, 2006). It is, therefore, possible that our participants are sub-optimal because they fail to take account of external noise, only weighting the sensory and prior information by the internal variability (i.e., error intrinsic to them) of the sensory cue. Keeping this is mind, we considered the predicted weights of a model that only takes into account the internal variability 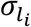 in using the sensory cue.

We performed a separate control experiment to see how good participants were at finding the centroids of dot clouds in the absence of prior information (see Appendix B for more details). From this control data, we could calculate observers’ internal noise as their responses were not subject to bias from the prior. For each participant, their internal variability 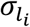 was calculated by taking the standard deviation of their errors from the centroid of the dots (error = dot centroid – response). The predicted weight on the sensory cue was then calculated as 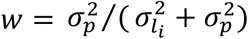. This equation was the same as the full Bayesian model, the only difference being that the external SD of the likelihood (as defined by the experimenters) was substituted for the internal SD of the likelihood (measured in the control experiment). The variance of the prior was still included in the model.

We compared subjects’ weights (block 5, Experiment 1) with those predicted when only weighting by internal noise and found that they were significantly different for all likelihood and prior pairings (*p* < .01). Indeed, Figure 5 shows that the internal noise model still predicts less weight on the sensory cue than we see in our data (compare bars and dotted lines). This could reflect participants downweighing the prior because it is, in fact, subject to additional internal noise, stemming from a need to remember and recall the correct prior from memory. Even so, empirical weights were closer to the internal noise predictions, compared to those, predicted by the optimal strategy (with experimentally-controlled cue variance, dashed lines).

**Figure 5.**
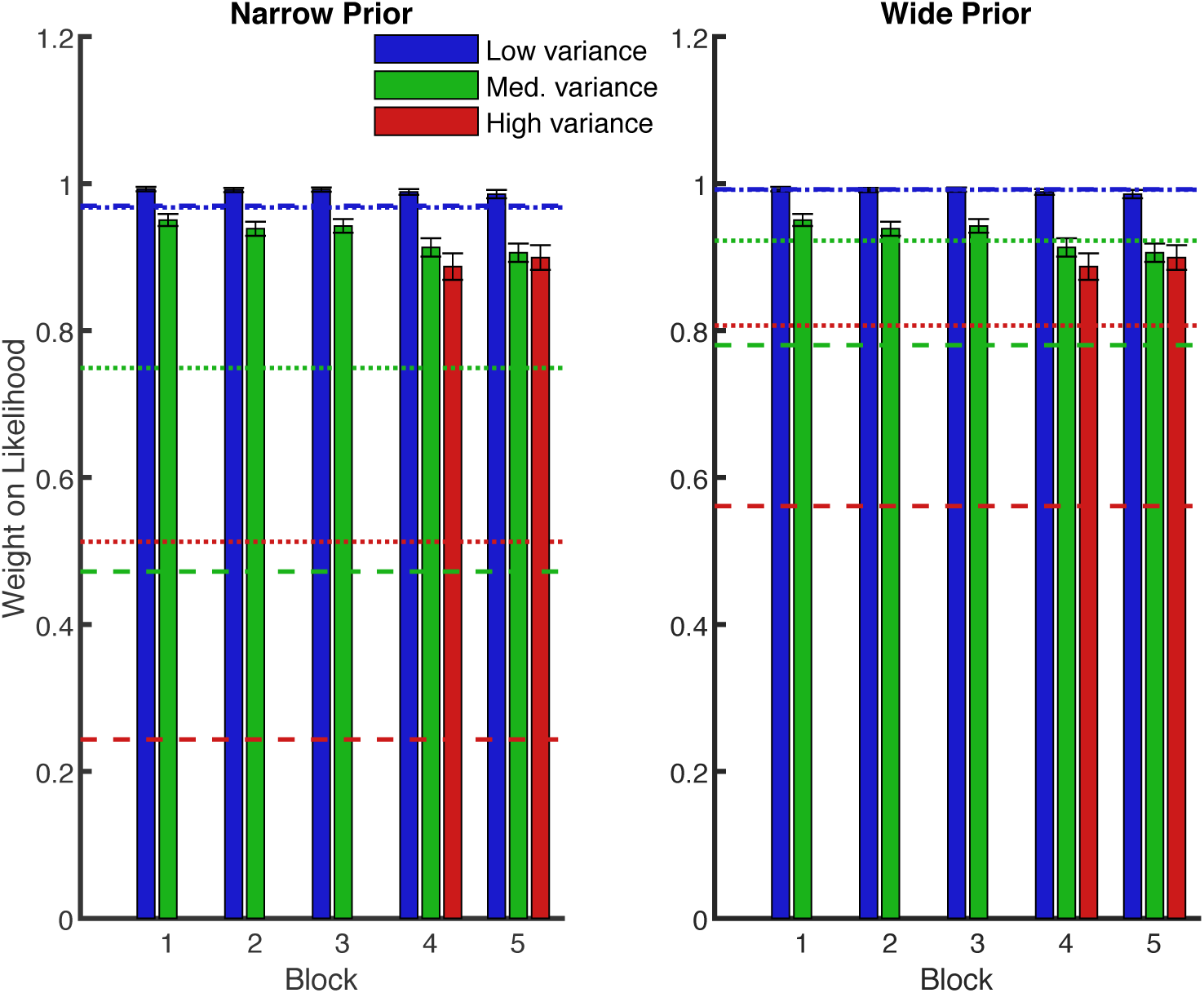
Mean weight placed on the likelihood information in each block of Experiment 1. Blue is the low-variance likelihood, green is the medium-variance likelihood, red is the high-variance likelihood. Dashed lines show optimal values. Dotted lines show predicted weights when only weighting by internal noise. Error bars are +/− 1 SEM.

In addition, we examined the predictions for an observer model that weights sensory information, according to overall variability in the sensory cue. When calculating the optimal predicted weights initially, we assumed that the optimal observer knew how reliable the dots (i.e., the likelihood) were, and could average them perfectly. Since participants will not be perfect at averaging dots, they will be more variable in using the sensory cue than the optimal observer. Therefore, the truly optimal thing to do is for participants to weight the sensory cue, according to their overall variability by taking into account both the variance of the dot distribution and the internal variability in estimating the average of the dots. Since the sensory cue is now less reliable (due to the added internal variability), we would expect participants to put less weight on it, and more weight on the prior.

We calculated the overall variability in using the sensory cue as:

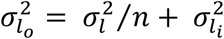

where 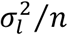 is the external noise in the sensory cue, and 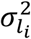 is the individual internal variability.

As is expected, Figure 6 shows that the predicted weights in this case were lower than the optimal weights (compare dotted and dashed lines) as participants are worse than the optimal observer in averaging the dots. They placed less weight on the sensory cue and more weight on the prior than the optimal observer. We also compared these predicted weights to subjects’ weights in the final block (5) in Experiment 1, and found that they were still significantly different from the empirical data when the variance of the likelihood was medium or high (irrespective of prior variance) and when the likelihood variance was low and the prior variance was narrow (all *p* < .001). No significant difference was observed when the likelihood variance was low, and the prior variance was wide (*p* = .79). This means that accounting for the added internal variability fails to explain our results as observers are placing more weight on the sensory cue than is optimal, not less.

**Figure 6.**
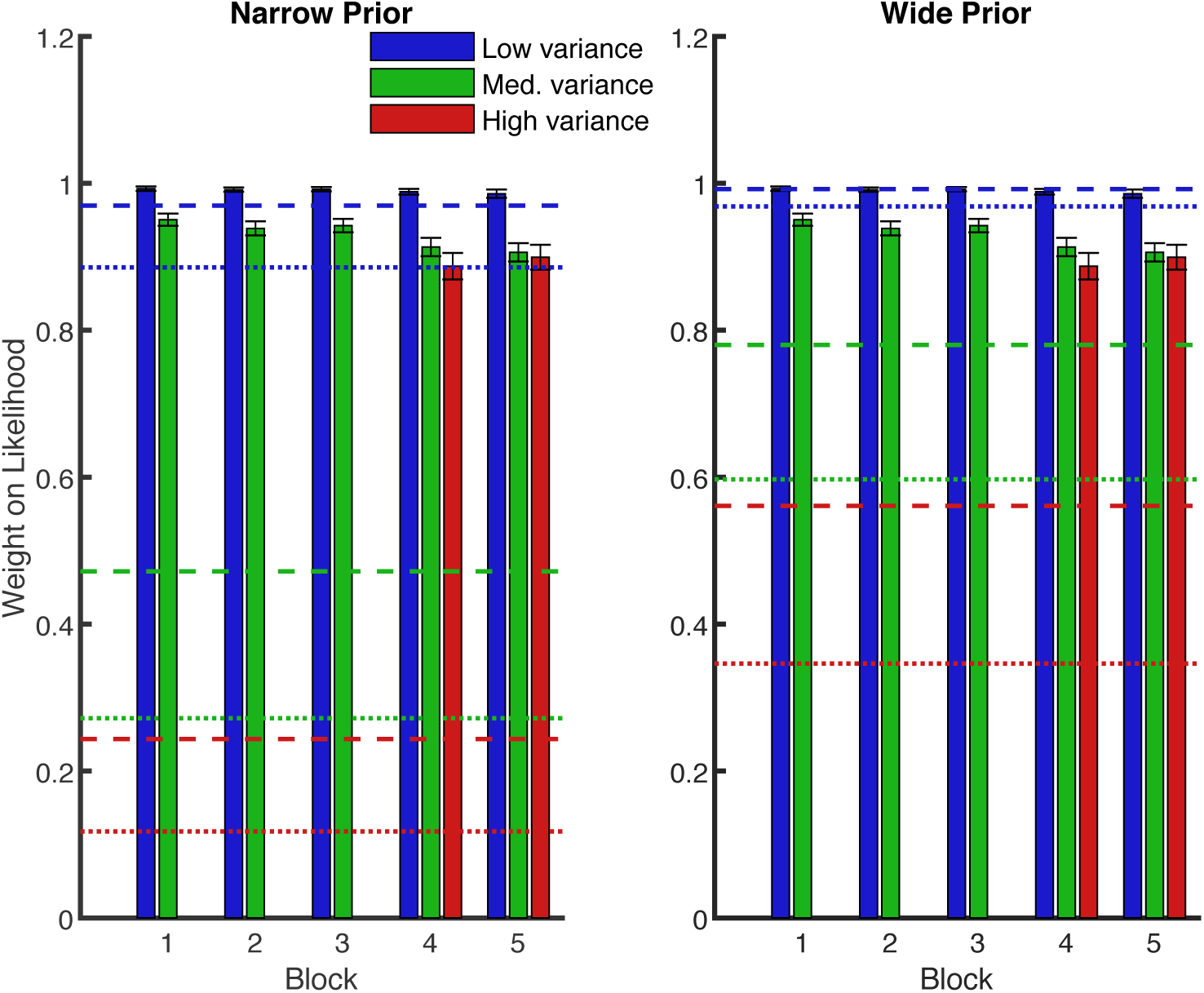
Mean weight placed on the likelihood information in each block of Experiment 1. Blue is the low-variance likelihood, green is the medium-variance likelihood, red is the high-variance likelihood. Dashed lines show optimal values. Dotted lines show predicted weights when overall variability in using the likelihood is taken into account. Error bars are +/− 1 SEM.

We compared the mean squared error (MSE) for each of the three models we tested: a) the original optimal model (using the experimentally imposed likelihood variance); b) the model with only the internal noise; and c) the model with the overall variability (including both the experimentally imposed likelihood variance and the internal noise). The internal noise model had the lowest MSE, which confirms that this model provides a better explanation for subjects’ behaviour than other models (see Figure 7).

**Figure 7.**
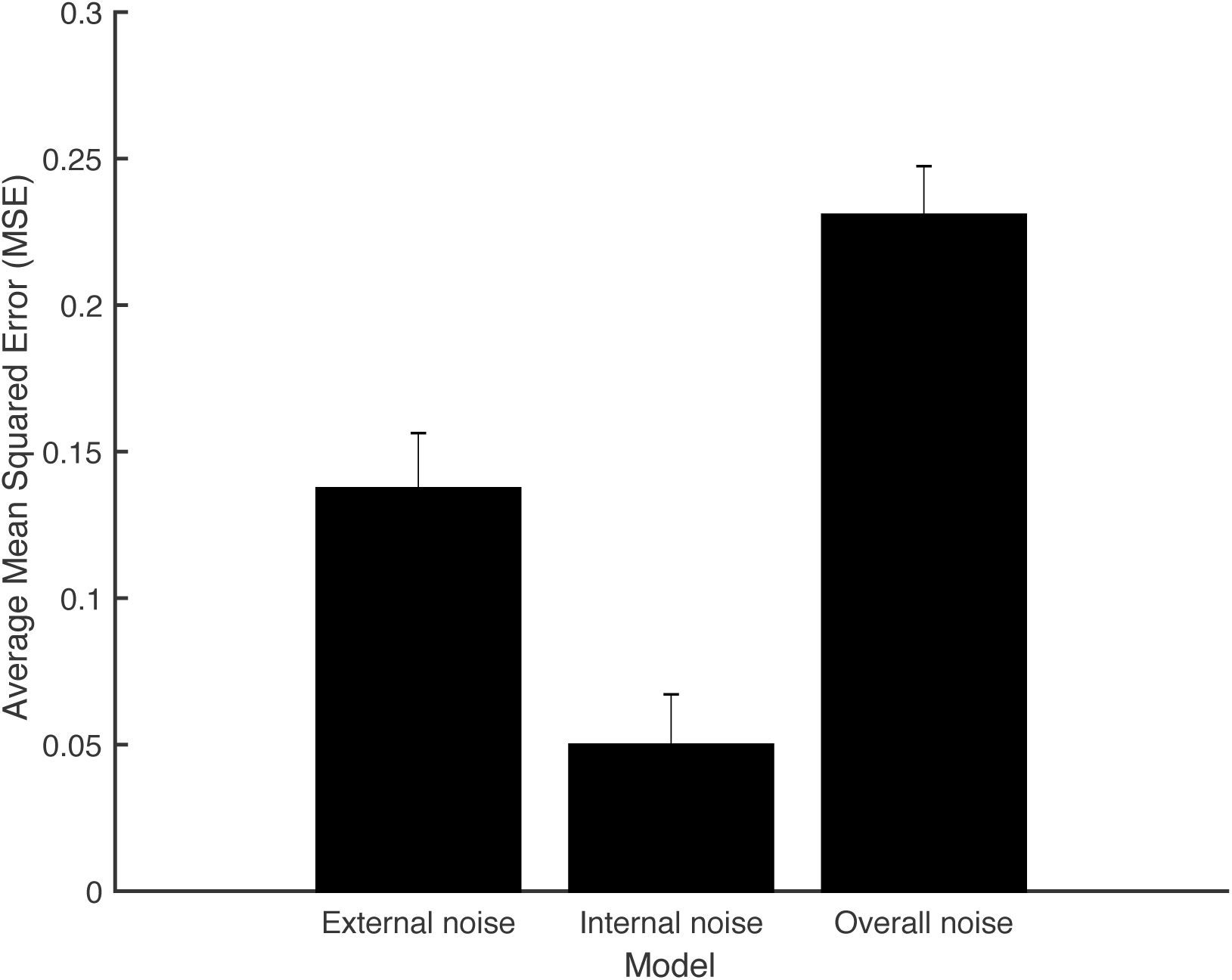
Average Mean Squared Error (MSE) for the external noise, internal noise and overall noise models. MSE calculations were based only on participants, for whom control data was available (N = 12; 6 had participated in Experiment 2 and 6 had participated in Experiment 3).

In summary, our data are best described by a model based on subjects’ internally generated noise, as opposed to either a model with the experimentally imposed likelihood variance, or a model that accounts for both the experimentally imposed likelihood variance and the internal noise.

## Discussion

We set out to test more strictly than before for Bayes-like combination of prior knowledge with sensory information in the context of a sensorimotor decision making task (Beierholm, Quartz, & Shams, 2009; Bejjanki et al., 2016; Berniker et al., 2010; Tassinari et al., 2006; Vilares et al., 2012) by adding transfer criteria to the task (Maloney & Mamassian, 2009).

In Experiment 1, we did this by investigating whether observers are able to learn the variances of two prior distributions, and instantly integrate this knowledge with a new level of sensory uncertainty added mid-way through a task. We found that observers placed more weight on the sensory cue (likelihood) when its variance was low and the variance of the prior was high; behaviour that is in broad agreement with the Bayesian prediction. However, we found only partial evidence of transfer. The weight placed on the high variance likelihood was not significantly lower than that placed on the medium variance likelihood, which is at odds with the prediction of transfer. Importantly, even though qualitatively, our participants behaved like Bayesian observers, their performance fell markedly short of optimal.

In two further experiments we asked: (1) how behavior would be affected by additional instructions, which can clarify whether this suboptimality stems from using the incorrect model structure of the task; (2) whether experiencing the high likelihood variance condition from the start of the experiment would lead to closer-to-optimal weighting of the prior and likelihood information. In the first of these two further experiments, Experiment 2, we found that subjects’ performance moved closer to optimal when they were (indirectly) instructed that the prior variances were different – possibly why we were able to detect significant evidence of transfer in the task. However, they were still significantly sub-optimal in multiple experimental conditions. Participants remained significantly sub-optimal in the final experiment (Experiment 3), when the need for transfer was removed (all trials types were present from the start of the task) and the experiment became a more direct replication of (Bejjanki et al., 2016).

### Suboptimal weighting of prior and likelihood information

We show that observers take uncertainty into account, giving more weight to the sensory cue as its variance decreases, a result that is consistent across all three experiments. Equally, for a Bayes-like observer, we expect to find that the weight on the sensory cue is higher as the prior variance increases, but we found a main effect of prior in Experiments 1 and 2 only, and not in Experiment 3. Moreover, while our manipulation to the instructions in Experiment 2 moved the weights placed on the likelihood closer to optimal, they were still significantly different to the optimal predictions.

To examine to what extent additional sensory variability in estimating the centres of dot-clouds could have affected predictions and performance, we ran a separate, control experiment (see Appendix B). This shows that observers are less efficient in their use of the likelihood information than an ideal observer: the variability of their responses is significantly larger than the true variability of the sensory cue in both the low and medium variance likelihood conditions. However, this fails to account for suboptimal performance: ideal weights for the likelihood that are computed using the measured likelihood variabilities in the control task are still significantly lower than those in the empirical data.

Suboptimal weighting of the prior and likelihood information may also be caused by incomplete or incorrect learning of the prior information. However, the prior-only trials suggest that the observers learn the means of the priors and distinguish between their variances at least in Experiments 1 and 2, if not Experiment 3 (under the assumption that standard deviations of subjects’ responses are related to the learnt prior variances). Suboptimal weighting of the prior could also be due to the use of an incorrect Bayesian generative model (causal structure) by subjects, e.g. if they believe that the prior will change over trials then they should apply a smaller weight to the prior (could be conceptualized as a meta-prior or hyperprior in Bayesian terms, Gelman, Carlin, Stern, & Rubin, 2013). The fact that we found an effect of instructions imply that the causal structure assumed by subjects can indeed greatly influence behavior (Shams & Beierholm, 2010).

Other research groups have performed similar experiments but using only a single prior distribution (Acerbi, Vijayakumar, & Wolpert, 2014; Berniker et al., 2010; Chambers et al., 2018; Tassinari et al., 2006; Vilares et al., 2012). These studies also find deviations from the optimal predictions; however, the deviations can be accounted for by adding extra sources of inefficiency to the model that are due to motor errors, centroid calculation errors, and aim point (in reaching tasks) calculation errors (Tassinari et al., 2006). Moreover, when trials are blocked by prior condition, it has been shown that learning after a switch in the prior variance is slower when the prior variance decreases than when it increases, suggesting participants may perform optimally after further exposure to the task (Berniker et al., 2010).

We considered elements of the experimental design that could have resulted in suboptimal behavior in a task similar to others in the literature where performance was closer to optimal (e.g., Bejjanki et al., 2016). Firstly, describing the dots as the tentacles of an octopus may have caused participants to assume that another method of gaining an estimate from the dot colour, other than taking the mean horizontal position, was more appropriate in this task (de Gardelle & Summerfield, 2011; Van Den Berg & Ma, 2012). However, our analysis shows that participant responses are not better predicted by the median, mid-range, or robust average, than they are by the mean. Secondly, our correction of the dot positions so that their SD on each trial was equal to the true likelihood SD may have influenced participant’s inferred reliability for the likelihood. However, an observer who computes the reliability for the likelihood trial by trial, by taking the dot cloud SD, would infer that the likelihood was less reliable 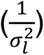 as a cue to true location than the centroid of the dots would be for an observer who could perfectly calculate the mean of the dots 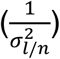. This would lead to an observer placing less weight on the likelihood than the ideal observer. Participants in our experiment place more weight on the likelihood than the ideal observer, so this is not the source of suboptimality in our experiment.

Finally, whilst the true likelihood and prior reliabilities used in our task were matched to those in Bejjanki et al. (2016) observers may have perceived the cue (dots) as more reliable than it actually was, which in turn would result in more weight placed on the cue than in previous studies (Bejjanki et al., 2016; Vilares & Kording, 2011).

It is possible that subjects did not experience enough trials of each prior and likelihood uncertainty to reach optimal performance, and indeed, we find evidence of decreasing weights on the likelihood with increasing task exposure in both Experiments 1 and 2 (main effect of block, although not the rise from block 4 to 5 in Experiment 2). Crucially, however, our participants experienced more trials per prior than in Bejjanki et al. (2016) (750, compared to 600) where weights were closer to optimal, ruling out the possibility that observers did not experience enough trials to learn the complex features of the distributions.

The result that our participants’ performance was so different -in terms of level of sub-optimality-compared to Bejjanki et al. (2016) might be explained by a difference in instructions. Specifically, their instructions have a social element that ours do not, i.e., in their task, participants were instructed to interpret the likelihood dots as “locations that other people have previously guessed the bucket is located at” (vs. tentacles of the octopus in ours). This means that in Bejjanki et al. (2016) participants would have to take into account how accurate they think other people’s guesses are. If we assume that people give lower weight to information that is allegedly based on other people’s guesses, this might explain why observers in Bejjanki et al. (2016) generally weighted the likelihood less than in our experiments (Martino, Bobadilla-suarez, Nouguchi, Sharot, & Love, 2017). Another aspect about the instructions that is worth mentioning here is that, perhaps, our participants are more likely to assume that the body of an octopus is in the center of its tentacles, compared to previous guesses of other participants (Bejjanki et al., 2016) or splashes from a coin (Tassinari et al., 2006; Vilares et al., 2012). However, had this been the case, we would have expected participants’ responses to be better predicted by another estimate, such as the robust average, than the mean of the dots, and we found no evidence of this in the data.

Another explanation is that observers were being “optimally lazy”; that is, they deviated from optimal performance in a way that had minimal effects on their expected reward (Acerbi, Vijayakumar, & Wolpert, 2017). In this case, we would expect obtained reward to match well with the predictions of the optimal Bayesian model; instead, the predicted reward resulting from optimally combing sensory and prior information was higher than that obtained by our observers – particularly when the variance of the prior was narrow (Appendix C). Therefore, we have no reason to believe that the suboptimal behaviour we observed in our task was due to our participants being “optimally lazy” (Acerbi et al., 2017).

Nevertheless, we could show that the suboptimal behaviour in our task can be best explained by assuming that participants were weighting sensory information (relative to prior information) only according to the internal variability in using the cue, ignoring external noise. It is, thus, interesting to consider it as one potential explanation, on the computational level, for the deviations from optimal consistently reported in similar studies on combination of sensory and prior information (Bejjanki et al., 2016; Berniker et al., 2010; Sato & Kording, 2014; Tassinari et al., 2006; Vilares et al., 2012). However, we note that attending to internal noise may be easily mistaken for underweighting the total noise; future work could investigate the extent to which suboptimal behaviour is specifically linked to the use of internal variability, and not simply a general under-estimation of the total noise in the stimuli.

Note that the observer model, based on the internally generated noise, can still be considered “subjectively” optimal (fully Bayesian), in the sense that observers take into account and act according to their internal noise variability (Acerbi et al., 2014). This strategy looks sensible but is arguably not Bayes-optimal as an ideal observer has to take into account external sources of noise in addition to his or her own sensory uncertainty (Kersten, Mamassian, & Yuille, 2004; Knill & Richards, 1996).

### Evidence for Bayesian transfer

We found only partial evidence for transfer in Experiment 1, as there was no significant change in the weight placed on the likelihood between the medium and high likelihood conditions. In fact, subjects seemed to treat the high variance likelihood the same as the medium variance likelihood (that they had experience with), suggesting that observers did not adopt a statistically optimal Bayesian strategy. Nonetheless, performance did not improve with more trials, suggesting that subjects were not implementing a look-up table decision rule, either (Maloney & Mamassian, 2009). However, we note that in our data, observers placed much more weight on the medium and high likelihoods than is optimal. This means that the effect size of a change in likelihood is much less than was expected; thus, the observed lack of significant differences might simply be due to lack of statistical power in our analysis to detect such small effect sizes.

Why do we see more convincing evidence of transfer in the instructions task? Bayes-like computations demand considerable computational resources (e.g., working memory load, attentional focus); it is, therefore, reasonable to expect that if a task is sufficiently complex, and there is a lot to learn, subjects will start behaving sub-optimally. The impact of additional instructions in Experiment 2 may be to free up cognitive resources by providing subjects with (indirect) information about the variances of the two prior distributions at the start of the task (Ma, 2012; Ma & Huang, 2009).

Our findings do not allow us to clearly distinguish between the reinforcement-learning and Bayesian interpretations. We found that when we introduced a new level of (known) uncertainty to the likelihood, observers immediately changed how they used this new information in a way that is largely consistent with optimal predictions; this effect was significant in the second experiment, but not in the first. Thus, we note that this effect is not particularly robust as it depends on the experimental procedure used to measure it. Indeed, our findings demonstrate that whether this effect is observed in the first place is greatly affected by small changes in experimental layout (e.g., instructions, number of trials). The fact that we observe no learning during Experiment 3 (no main effect of block), coupled with the observation that the weight on the likelihood in the 4^th^ block of Experiment 3 were remarkably similar to those in the 4^th^ block of Experiments 1 and 2 makes a weak suggestion of a Bayesian interpretation. However, a stronger test of transfer would be if participants had received no feedback for the new level of uncertainty. We provided trial-by-trial feedback (true target position) to ensure that participants were able to learn and recall the correct prior distributions. Therefore, we cannot rule out the possibility that our participants used the feedback to directly learn a mapping between the high variance likelihood and each prior, instead of the distribution of locations.

Sato and Kording (2014) showed that subjects behave in a Bayes-optimal fashion in a sensorimotor estimation task, where they transferred their knowledge from the ‘learning phase’ to the prior in the testing phase in the absence of trial-to-trial feedback, suggesting that people did not learn a simple likelihood-prior mapping. This means that the features of our experiments set an approximate upper bound on learning; in other words, we can generally expect subjects’ performance to be less accurate when performance feedback is not provided.

Nevertheless, in order to meaningfully test whether observers can transfer probabilistic information across different conditions, an experiment where trial-by-trial feedback is limited, or excluded altogether, is needed. Hudson, Maloney, and Landy (2008) argued that providing only blocked performance feedback, for example, would prevent participants from using a “hill-climbing” strategy in the high variance likelihood condition (i.e., updating their estimates, based on the feedback from trial to trial). Alternatively, Acerbi, Vijayakumar, and Wolpert (2014) found that partial feedback (where participants are told whether they “hit” or “missed” the target, but the actual target position is not displayed) is sufficient to maintain participant engagement; however, no meaningful information can be extracted from the feedback, preventing participants from using it to better their performance. Future work could investigate how the removal of full performance feedback would affect behaviour in more complex scenarios.

### What are observers if not Bayesian?

Some studies suggest that BDT is generally a good descriptive model of people’s perceptual and motor performance, but quantitative comparison shows divergence from Bayes-optimal behaviour (Bejjanki et al., 2016; Zhou, Acerbi, & Ma, 2018), not unlike what we report in this study. These deviations from optimality may have arisen because rather than performing the complex computations that a typical Bayesian observer would do, observers draw on simpler non-Bayesian, perhaps even non-probabilistic, heuristics (Gigerenzer & Gaissmaier, 2011; Zhou et al., 2018). Laquitaine and Gardner (2018) developed a model that switched between the prior and sensory information, instead of combining the two, which was found to explain the data better than standard Bayesian models. The authors concluded that people can approximate an optimal Bayesian observer by using a switching heuristic that forgoes multiplying prior and sensory likelihood. In another study, Norton, Acerbi, Ma, and Landy (2018) compared subjects’ behaviour to the ‘optimal’ strategy, and well as several other heuristic models. The model fit showed that participants consistently computed the probability of a stimulus as belonging to one of two categories as a weighted average of the previous category types, giving more weight to those seen more recently; subjects’ responses also showed a bias towards seeing each prior category equally often (i.e., with equal probability). We note that a Reinforcement-Learning (RL) model was also tested, where participants could simply update the decision criterion after making an error with no assumptions about probability; no participant was best fit by the RL model. This suggests that observers are, in fact, probabilistic, i.e., take into account probabilities, though not necessarily in the optimal way; instead, they seem to resort to heuristic strategies. However, future work should explore which, if any, of these models can capture the behaviour on this type of complex localisation task.

## Acknowledgements

This work was supported by research grants from the Leverhulme Trust (RPG-2017-097; Marko Nardini, PI) and North East Doctoral Training Centre (ES/J500082/1; Reneta Kiryakova). We would like to thank Dr. James Negen for helpful comments on the manuscript.

## Appendix A

### Instructions in Experiment 1

“We will ask you to play an “octopus” game!

Imagine that you are on a boat and there are 2 octopuses you are trying to find: one is white, and the other one is black. The white octopus has square tentacles and the black octopus has circular tentacles. The 2 octopuses live in different parts of the sea. Sometimes the octopuses will show their tentacles and at other times they will hide at the bottom of the sea.

Your job is to try and figure out where the octopus is!

Once you decide on a location, you can click on the green square (your fishing net), at which point you will see a red dot, which shows you the true location of the octopus on that trial. If the red dot is inside the net, then you correctly guessed the location of the octopus and you get a point!”

### Instructions in Experiment 2

“We will ask you to play an “octopus” game!

Imagine that you are on a boat and there are 2 octopuses you are trying to find: one is white, and the other one is black. The white octopus has square tentacles and the black octopus has circular tentacles. The 2 octopuses live in different parts of the sea. Sometimes the octopuses will show their tentacles and at other times they will hide at the bottom of the sea.

**[It is important to remember that one of the octopuses tends to stay in a particular area, whereas the other one moves quite a bit!]**

Your job is to try and figure out where the octopus is!

Once you decide on a location, you can click on the green square (your fishing net), at which point you will see a red dot, which shows you the true location of the octopus on that trial. If the red dot is inside the net, then you correctly guessed the location of the octopus and you get a point!”

## Appendix B

### Control Experiment: Likelihood-only task

Participants consistently performed sub-optimally across *all* of our experiments. However, when we calculated the optimal weight on the likelihood, we did so under the assumption that people know the true values of the reliability of the sensory cue (i.e., the likelihood). As Sato and Kording (2014) point out, this is clearly not always the case: in fact, in order to perform optimally on our tasks, observers may need to learn about their likelihood variability, as well as prior variability. We, therefore, separately assessed any sensory noise that participants may have had in judging the centroid of the set of dots. If we find that subjects’ estimates of the reliability of the likelihood differ from the true values, this would mean that subjects were using incorrect parameters for the task, which may have led to suboptimal performance. We then recomputed the optimal weights based on errors in observers’ estimates of centroid location; we could, therefore, test whether subjects were, in fact, near-optimal, when their own sensory variability was taken into account.

### Methods

Subjects (*N* = 26; 6 had participated in Experiment 2, 6 had participated in Experiment 3, and the rest had not completed any of the above tasks) were instructed to estimate the centroid of eight dots for different likelihood widths. True locations were drawn from a uniform distribution across the screen (no prior). There were 90 trials overall, with 30 trials of each likelihood width interleaved in a random order. No feedback was given.

For each participant, their error on each trial was calculated by taking the difference between the response and true location for that trial (error = response – true). Their variable error for each likelihood condition was calculated as the standard deviation of the errors. Outliers were excluded prior to calculating the variable error in the same way as described previously.

### Results and Discussion of Control Experiment

Participants were significantly worse than ideal (variable error was greater than the true standard deviation of the likelihoods) in the low (*t*(25) = 7.45, *p* < .001) and medium (*t*(25) = 3.80, *p* < .001), but not the high (*t*(25) = 1.48, *p* = .151) variance likelihood conditions (Figure 8).

**Figure 8.**
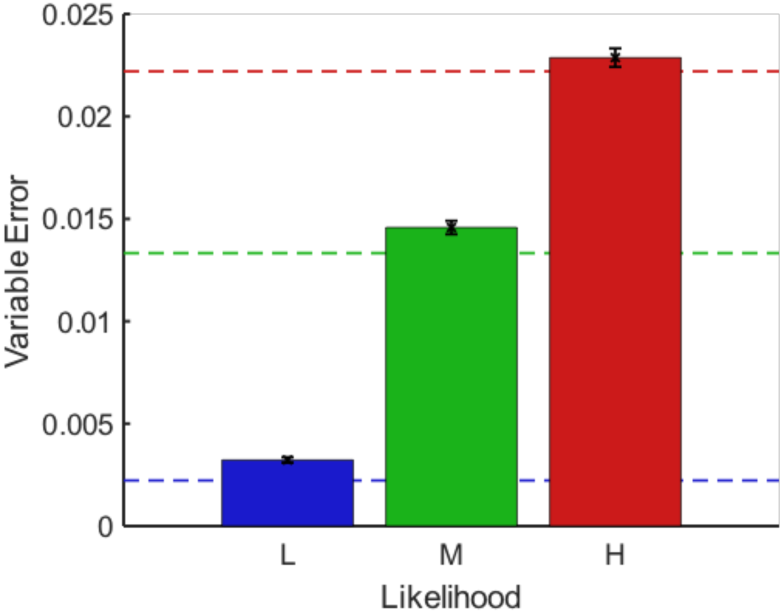
Variable error for each likelihood condition in the likelihood-only task. The dashed lines show the true standard deviations of the likelihood in each case (ceiling performance).

This suggests that our optimal predictions place too much weight on the likelihood, as they were calculated based only on the external variability of the sensory cue and failed to also incorporate the added variability from observer’s inability to perfectly calculate the dot centroids. We, therefore, recomputed the ideal weight for the likelihood, this time using the measured likelihood variances in the control experiment; we reasoned that this calculation would give us an optimal prediction that better matches our subjects’ performance. Our estimates of the likelihood variance increased by 16.66% for the low, 2.96% for the medium and 5.26% for the high likelihood. With such large differences between the true and estimated likelihood variances, we expected that the re-calculated optimal predictions (based on subjects’ estimates) will be closer to the observers’ data, compared to those based on the true likelihood parameters. We compared these optimal values to subjects’ weights in the final block (5) in Experiment 1, and found that they were still significantly different from the empirical data when the variance of the likelihood was high or medium, irrespective of prior variance (all *p* < .001) (see Figure 9). No significant differences were observed when the prior variance was wide and the likelihood variance was low (*p* = .765).

**Figure 9.**
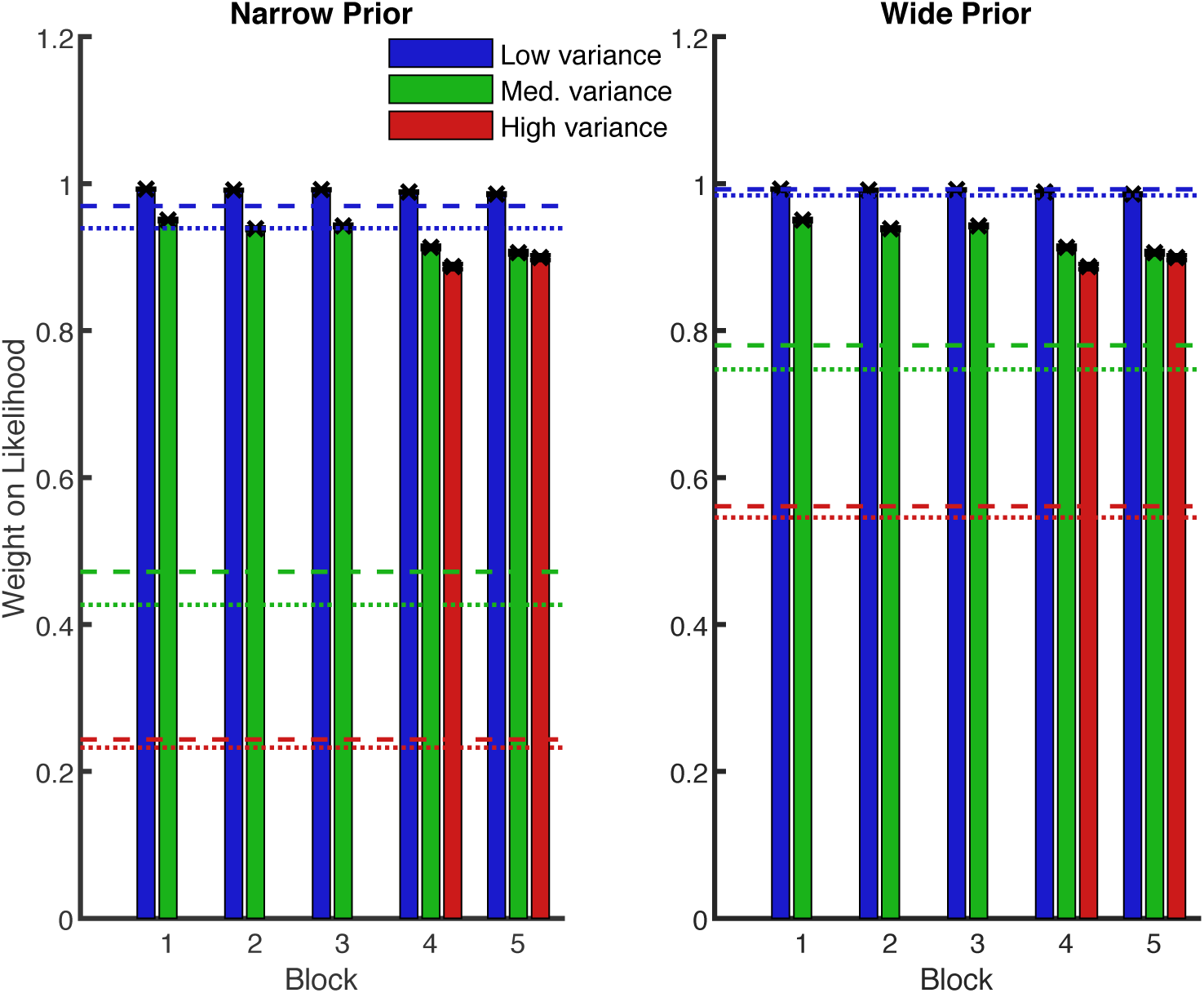
Mean weight placed on the likelihood information in each block of Experiment 1. Blue is low variance likelihood, green is medium variance likelihood, red is high variance likelihood. Dashed lines show optimal values. Dotted lines show optimal values, computed using measured likelihood variances in the control experiment. Error bars are +/− 1 SEM.

This pattern of results was surprisingly similar to the one we found when using the predictions of the optimal Bayesian observer, so this analysis did not affect our conclusions on the observers’ suboptimal behaviour. In particular, sensory noise in determining the centroid of the “likelihood dots” does not play a major role in explaining subjects’ sub-optimality in the task.

## Appendix C

The observed lack of statistically significant difference in cue weights does not necessarily imply a lack of substantial difference in terms of performance (points), as previous studies have shown that participants can be “optimally lazy” by deviating from optimal performance in a way that has minimal impact on overall expected score in a task (Acerbi et al., 2017). First off, we computed the optimal response variability 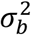 in using both the cue (overall likelihood variability 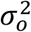) and the prior as

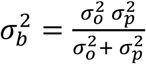

Since we are interested in the performance of the model in terms of reward, we then calculated expected gains by first computing the probability of catching an octopus on a given trial as

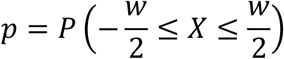

where *p* is the probability that a random draw *X* from a Gaussian distribution with mean *μ* (fixed at zero) and standard deviation 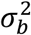 will fall within the “hit” distance from the true location, and that distance is half the width of the net 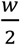. The probability of catching the octopus *p* is then multiplied by the number of trials (per trial type in a block) to calculate the expected number of points.

We then compared expected reward to the average reward earned by those participants who took part in the control experiment and either Experiment 2 or Experiment 3 (*N* = 12; in block 5 only), and found that optimal integration of the sensory cue and prior knowledge (according to participants’ overall noise in using the cue) resulted in an expected reward that was higher than what our participants achieved, but only when the variance of the prior was narrow; when the prior variance was wide, they matched quite well; see Figure 10. This result is particularly challenging for the notion that people may be “optimally lazy”, as this case would result in predicted and obtained reward values being equal. It can be seen that contrary to these predictions, our observers were clearly worse than the optimal observer, and could earn more points when the variance of the prior was narrow; it is, therefore, unlikely that their suboptimal performance could be explained by them being “optimally lazy” (Acerbi et al., 2017).

**Figure 10.**
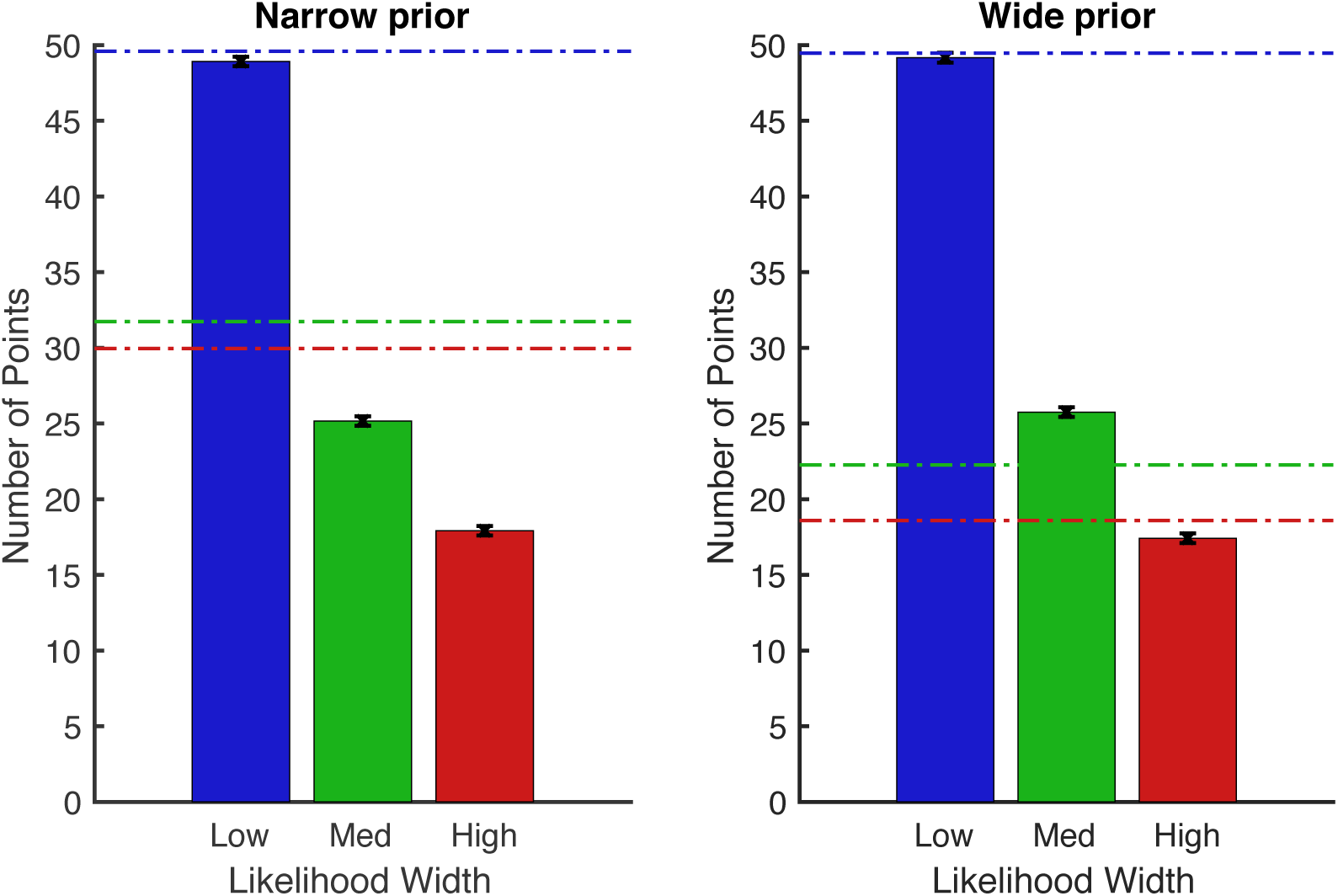
Mean number of points earned in Block 5 for participants who took part in Experiment 2 or Experiment 3 and the control task (*N* = 12). Blue is low variance likelihood, green is medium variance likelihood, red is high variance likelihood. Dot-dashed lines show optimal reward values, taking into account participants’ overall noise. Error bars are +/− 1 SEM.

